# PPGLomics: An Interactive Platform for Pheochromocytoma and Paraganglioma Transcriptomics

**DOI:** 10.64898/2026.01.29.702561

**Authors:** Hussam Alkaissi, Catherine M. Gordon, Karel Pacak

## Abstract

Pheochromocytoma and paraganglioma (PPGL) are rare neuroendocrine tumors with unique biological behavior and remarkably high heritability, yet dedicated bioinformatics resources for these diagnoses remain limited. Existing cancer multi-omics platforms are pan-cancer in scope, often lacking the disease-specific annotations, granularity, and cross-database harmonization required for meaningful stratification and hypothesis generation. Here we introduce PPGLomics, an interactive web-based platform designed for comprehensive PPGL transcriptomics analysis. PPGLomics v1.0 integrates two major datasets, the TCGA-PCPG cohort (n=160) spanning multiple molecular subtypes, and the A5 consortium SDHB cohort (n=91) with detailed clinicopathological and molecular annotations. The platform provides basic and clinical scientists, as well as a broad range of healthcare professionals, with tools for differential expression analysis, correlation analysis, survival analysis, and visualization, including boxplots, heatmaps, volcano plots, and Kaplan-Meier survival plots, enabling exploration of gene expression patterns across PPGL subtypes without requiring bioinformatics expertise. PPGLomics v1.0 is freely available at https://alkaissilab.shinyapps.io/PPGLomics.

**Graphical Abstract:** 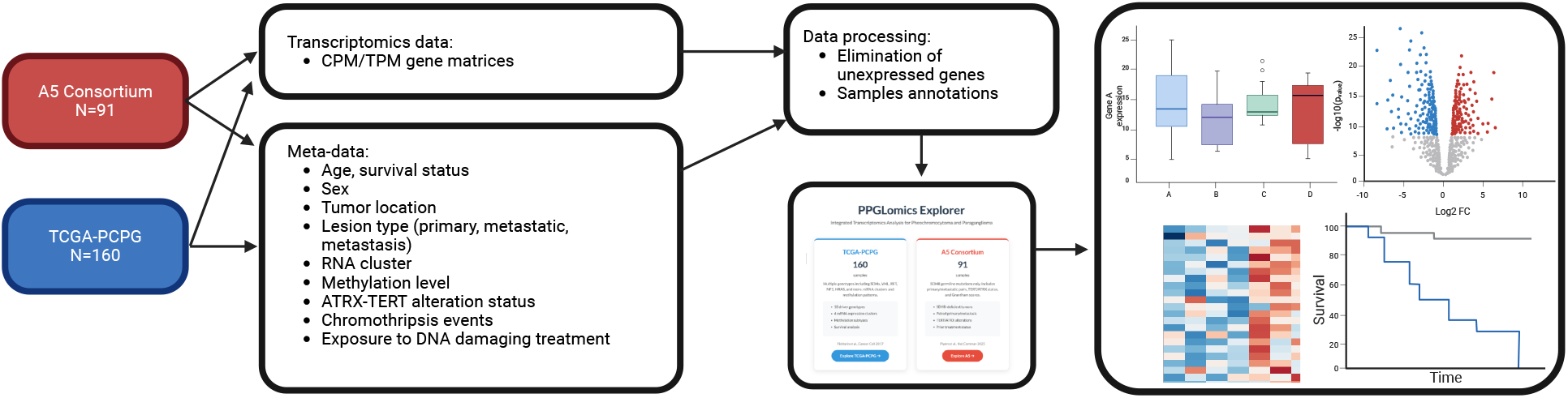

## Introduction

The publication of the human genome at the turn of the century required substantial financial investment, international collaboration, and a coordination between public and private sectors (1). In the following 25 years, sequencing costs have decreased by several orders of magnitude while throughput and accuracy have dramatically improved (2). The speed at which results are available has also dramatically increased. Whole-genome sequencing can now be performed and interpreted within a single day, shifting the primary bottleneck in genomics from data acquisition to data analysis and interpretation, all tightly linked to the individual patient scenario (2, 3).

Large-scale cancer genomics initiatives, most notably The Cancer Genome Atlas (TCGA), have generated multi-omics data across different cancer types, creating unprecedented opportunities for hypothesis-generation and discoveries (4). To make these data accessible to biologists and clinicians, several user-friendly web portals have been developed by large bioinformatics teams, such as cBioPortal, GEPIA, UALCAN, OncoDB, and UCSC Xena (5-10). These tools have democratized access to cancer genomics data, and facilitated cross-cancer multi-omics comparisons, enabling researchers and clinicians without bioinformatics expertise to conduct sophisticated genomic analyses. GEPIA alone processed over 400,000 analysis requests within two years of its launch (6).

Existing platforms are pan-cancer by design and are of limited utility in rare cancers research that require disease-specific context and annotations. Pheochromocytoma and paraganglioma (PPGL) exemplify this challenge. Paragangliomas are tumor of autonomic ganglia (sympathetic or parasympathetic), many of which are hormonally active, secreting large amounts of catecholamines. Anatomic location of paragangliomas infer some of their behavior. For example, head and neck paragangliomas are of parasympathetic origin, and are less likely to metastasize and more likely to follow an indolent course, while abdominal paragangliomas are of sympathetic origin with higher rates of metastasis and aggressive behavior (11, 12). These neuroendocrine tumors differ fundamentally from most malignancies, as they are slow growing yet can metastasize, often many years after the initial diagnosis (13). There is no definitive histopathological distinction between benign and malignant tumors, and virtually all tumors carry metastatic potential (14). Furthermore, PPGLs are among the most heritable cancers, with germline pathogenic variants identified in over 30% of patients and somatic drivers in approximately 40% (15-18).

The biological and clinical heterogeneity of PPGLs is currently largely determined by genotype. For example, tumors driven by mutations in genes encoding enzymes of Krebs cycle (e.g., succinate dehydrogenase subunit B, *SDHB*), extra-adrenal location, larger size, and tumors harboring *ATRX/TERT* alterations exhibit higher metastatic rates (13, 19). Meaningful PPGL transcriptomics analysis therefore requires genotype-specific stratification and disease-relevant annotations, including tumor location, size, ATRX/TERT status, that are usually unavailable in existing pan-cancer portals. An additional challenge is the scarcity of large cohorts in rare cancer research. The two largest publicly available PPGL datasets are the TCGA-PCPG cohort (n= 179, 19 excluded after pan-cancer quality control harmonization), spanning multiple genotypes with both primary and metastatic tumors, and the A5 Consortium cohort (n=91), comprising *SDHB*-driven tumors across anatomic locations with primary and metastatic tumors (16, 19).

To address this gap, we have developed PPGLomics v1.0, a free interactive web-based platform dedicated to PPGL transcriptomics analysis, with disease-specific annotations, providing a specialized tool for hypothesis generation and data exploration (Link: https://alkaissilab.shinyapps.io/PPGLomics).

## Methods

### Data Sources and Processing

PPGLomics v1.0 integrates two publicly available PPGL transcriptomic datasets, the TCGA-PCPG and the A5 Consortium (16, 20). RNA sequencing data for both cohorts were downloaded from cBioPortal; clinical and molecular annotations were obtained from the original publications.

The TCGA-PCPG dataset comprises 160 pheochromocytoma and paraganglioma samples (of 179 original cases following pan-cancer quality control) spanning multiple driver genotypes, including *SDHx, VHL, RET, EPAS1, EGLN1, HRAS, NF1, MAX, TMEM127, MAML3* fusions, *CSDE1*, and sporadic cases. Gene expression values are provided as log_2_-transformed transcripts per million (TPM). Metadata annotations include driver genotype and RNA expression cluster (pseudohypoxia, kinase signaling, Wnt-altered, and cortical admixture), lesion type (primary tumor, metastatic tumor), overall and progression-free survival, as reported by Fishbein et al. (16). Expression data were filtered to the 15,000 most variable genes to optimize web application performance, excluding genes with very low expression (genes with maximum expression < 2 log_2_ TPM across samples were excluded).

The A5 Consortium dataset consists of 91 PPGL samples (of 94 original cases following pan-cancer quality control), all harboring germline *SDHB* pathogenic variants (truncating, missense, frameshift, deletions, or intronic variants), as reported by Flynn et al. (20). Annotations include tumor anatomical location, lesion type (primary tumor, metastatic tumor, primary tumor from patients who subsequently developed metastases, and paired primary-metastasis cases), biochemical phenotype, tumor size, mutation type and position, *ATRX/TERT* alteration status, chromothripsis events, overall survival and metastasis-free survival. The initial dataset of over 42,000 genes was filtered to remove genes with very low expression (maximum expression < 2 log_2_ CPM). We included protein-coding genes, as well as differentially expressed novel loci (e.g., metastatic vs. primary tumors), including microRNAs and lncRNAs, to facilitate hypothesis generation and identification of new targets in PPGL biology. This filtering removed 43% of the initial expression matrix, yielding a final curated dataset of 23,928 genes.

### Web Application Implementation

PPGLomics was developed as an interactive web application using R Shiny package. The application implements: (1) single-gene expression analysis with boxplots and Kruskal-Wallis testing for nonparametric multi-group comparisons; (2) hierarchical clustering heatmaps with Z-score normalization; (3) pairwise gene correlation analysis using Spearman or Pearson correlation methods; (4) genome-wide similar gene identification via vectorized correlation; (5) differential expression analysis using Wilcoxon rank-sum tests with Benjamini-Hochberg correction, visualized as interactive volcano plots; (6) Kaplan-Meier survival analysis with log-rank testing; and (7) OncoPrint visualization of driver alterations. An example is shown in Figure 1.

**Figure 1.**
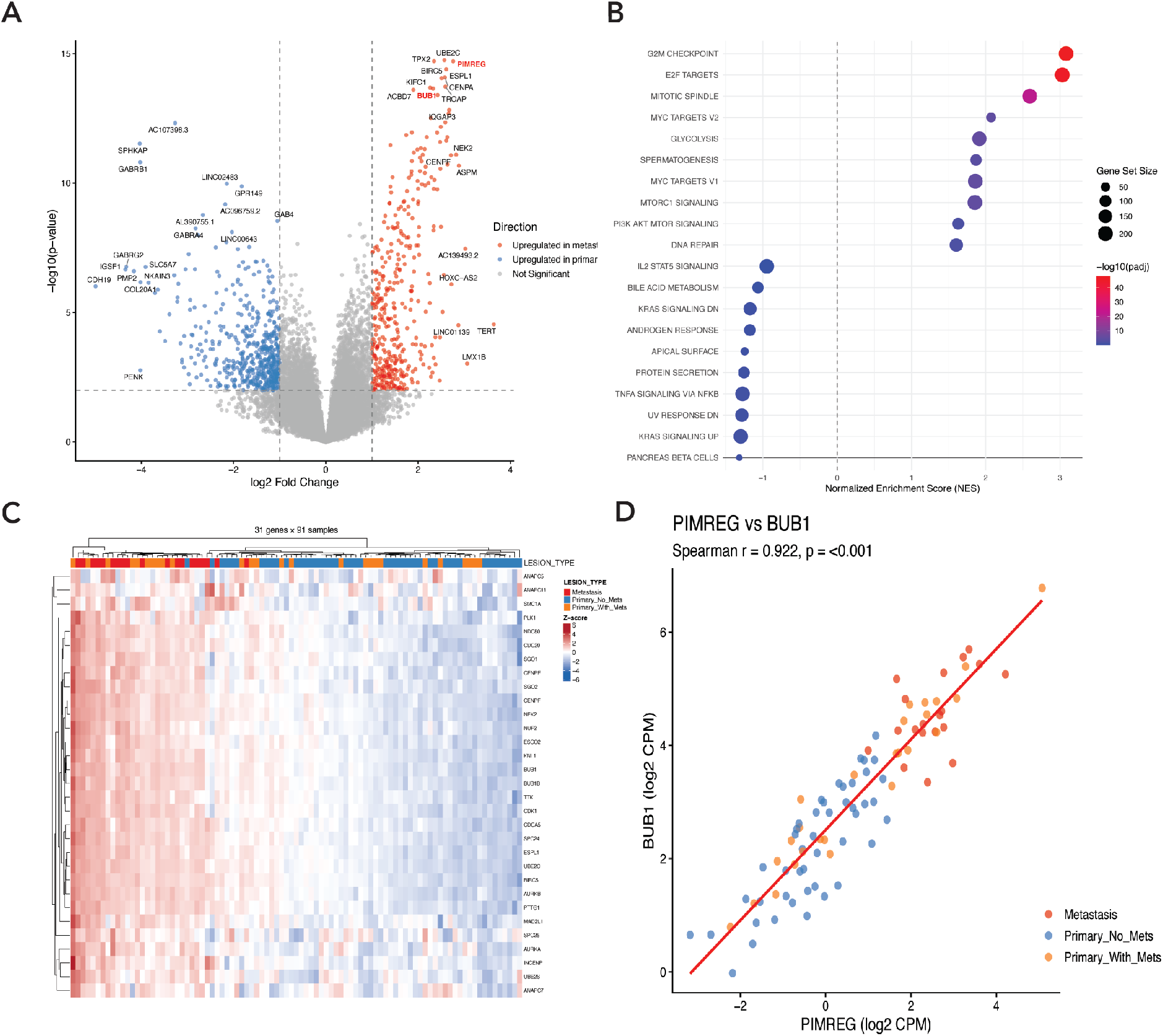
Mitotic spindle gene expression signature distinguishes metastatic *SDHB* PPGLs from primary *SDHB* PPGLs. (A) Volcano plot of differentially expressed genes between metastatic (n=19) and non-metastatic primary PPGL (n=46). Red and blue points indicate genes significantly upregulated in metastases vs non-metastatic primary tumors, respectively. Dashed lines denote significance thresholds (adjusted p < 0.05, log2 fold change > 1). (B) Gene set enrichment analysis (GSEA) demonstrating enrichment of cell cycle and mitotic pathways in metastatic samples. (C) Heatmap of selected 31 mitotic spindle-related genes across 91 samples. Column annotations indicate lesion type: non-metastatic primary (blue), primary with subsequent metastasis (yellow), and metastasis (red). Expression values are z-score normalized. (D) Correlation between *PIMREG* and *BUB1* expression (Spearman R = 0.92, p < 0.001).

Core R packages include Shiny for the web framework, ggplot2 and plotly for static and interactive visualizations, ComplexHeatmap for heatmaps and mutation landscapes, survival and survminer for Kaplan-Meier analysis, and matrixTests for vectorized statistical testing. The application was developed with AI assistance (Anthropic Claude) for code generation, with iterative testing and domain-specific refinement by the investigator.

### Case Study: Mitotic Spindle Gene Expression Signature in Metastatic PPGL

To identify transcriptomic signature associated with metastatic progression in *SDHB*-driven PPGL, we performed differential gene expression analysis comparing primary tumors without metastasis (n=46) to metastatic lesions (n=19) using the A5 Consortium dataset. This analysis revealed striking enrichment of genes involved in cell cycle regulation (E2F targets, G2M checkpoint), mitotic spindle assembly, and chromosome segregation among the most significantly upregulated transcripts in metastatic *SDHB* PPGLs (Fig. 2A-B). A focused analysis of 31 mitotic spindle-related genes demonstrated coordinated upregulation in metastatic samples, with notably elevated expression also observed in a subset of primary *SDHB* tumors that subsequently developed metastases (Fig. 2C). Among these genes, *PIMREG* (PICALM Interacting Mitotic Regulator) and *BUB1* (Budding Uninhibited by Benzimidazoles 1) emerged as potential markers of metastatic potential. Expression of these two genes was strongly correlated (Spearman R= 0.92, p < 0.001; Fig. 2D), suggesting coordinated activation of the mitotic program.

**Figure 2.**
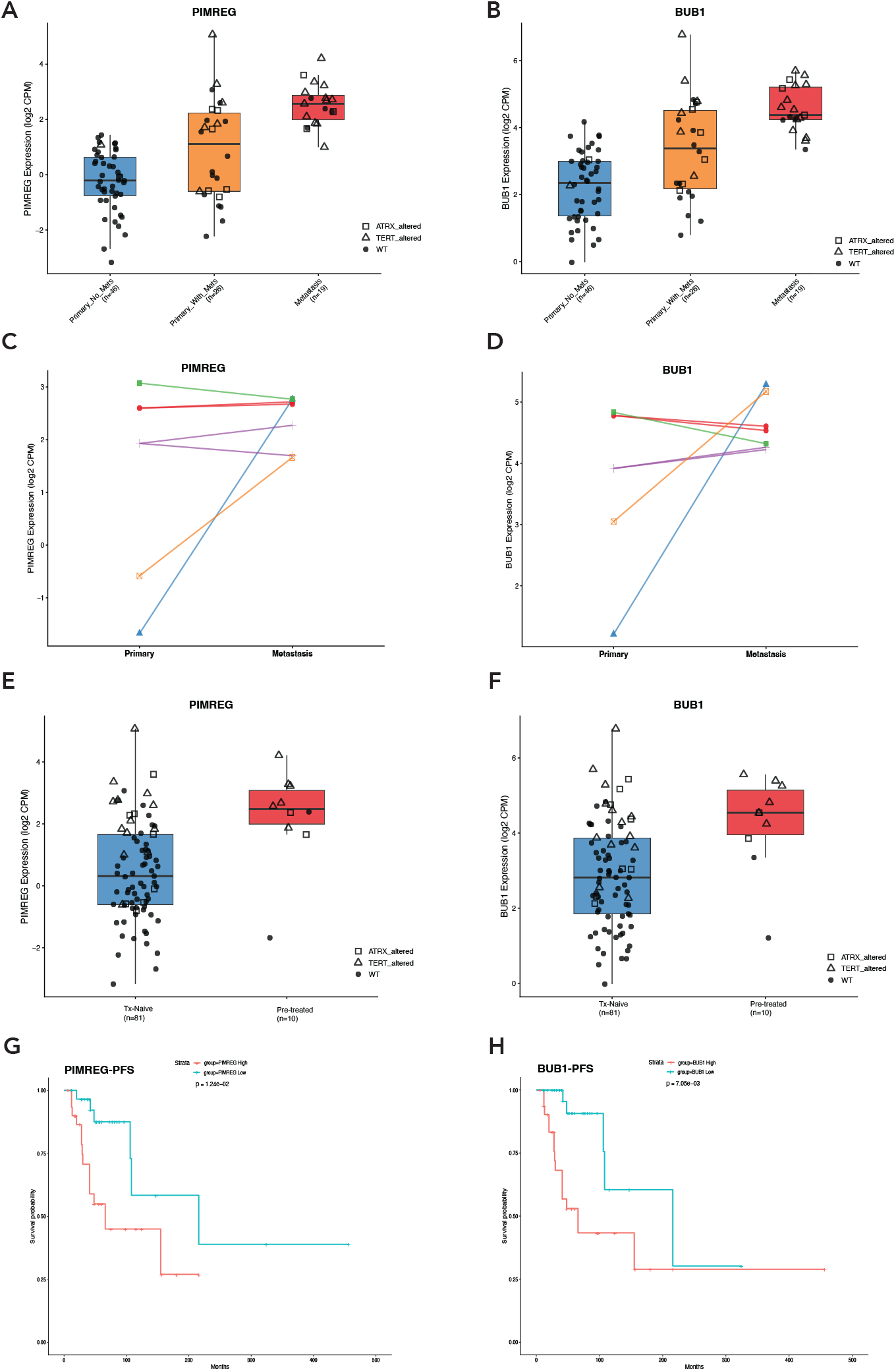
PIMREG and BUB1 expression is associated with metastatic progression and treatment history. (A-B) *PIMREG* and *BUB1* expression (log2 CPM) across *SDHB* lesion types. Open triangles and squares denote *ATRX/TERT* alterations, respectively (Kruskal-Wallis p = 5.87×10^−9^ and 4.94×10^−9^). (C-D) Paired analysis of matched primary-metastatic tumors; lines connect samples from the same patient. (E-F) Expression by prior treatment with DNA-damaging agents such as chemotherapy or radiotherapy. (G-H) Kaplan-Meier analysis of metastasis-free survival stratified by *PIMREG* and *BUB1* expression.

Both *PIMREG* and *BUB1* displayed stepwise expression across disease states, with lowest expression in primary tumors without metastasis (median log_2_ CPM = −0.21 and 2.35, respectively), intermediate in primary tumors that later metastasized (median = 1.11 and 3.38), and highest in metastatic lesions (median = 2.57 and 4.37; Kruskal-Wallis p = 5.87×10^−9^ and 4.94×10^−9^ respectively, Fig. 3A–B). Analysis of paired primary and metastatic *SDHB* samples from the same patients confirmed this pattern, with *PIMREG* and *BUB1* consistently upregulated upon metastatic progression in the majority of cases (Fig. 3E-F). Notably, expression of both genes was higher in tumors from patients previously treated with DNA-damaging agents such as CVD chemotherapy or ^131^I-MIBG (Fig. 3C-D), consistent with prior observations that alkylating agent exposure increases tumor mutational burden and may select aggressive clones (21).

Importantly, high *PIMREG* and *BUB1* expression in PPGLs was associated with significantly reduced metastasis-free survival (Fig. 3G-H).

PIMREG and BUB1 are mitotic regulators; PIMREG promotes the metaphase-to-anaphase transition while BUB1 is a central component of the spindle assembly checkpoint that restrains this transition until all chromosomes achieve proper bi-orientation (22-25). Beyond its canonical mitotic functions, PIMREG drives tumor aggressiveness through NF-κB activation and TWIST1 stabilization, promoting epithelial-mesenchymal transition (23). The highest PIMREG expression is observed in glioblastoma, where it contributes to temozolomide resistance (22).

BUB1 is a master coordinator of kinetochore assembly and inhibits the anaphase-promoting complex (APC/C), thereby preventing premature sister chromatid separation (24). BUB1 exhibits a tumor suppressor paradox where partial loss-of-function promotes aneuploidy and tumorigenesis, while complete inactivation is lethal to cells, a vulnerability that may be therapeutically exploitable (24). The strong co-expression of PIMREG and BUB1 in metastatic PPGL may reflect a coordinated transcriptional activation of the mitotic program enabling sustained proliferation despite the metabolic stress inherent to SDH-deficient state. PIMREG and BUB1 may serve as prognostic biomarkers for metastatic potential in *SDHB*-related PPGL. The dependence of metastatic PPGLs on mitotic checkpoint machinery may present a therapeutic opportunity for agents such as Aurora kinase A inhibitors, and CDK4/6 (26, 27).

## Discussion

PPGLomics v1.0 addresses the unmet need for disease-specific transcriptomics analysis tools in PPGL research. While pan-cancer portals such as cBioPortal and GEPIA provide very valuable resources for common malignancies, their utility for rare cancers is limited by the absence of disease-relevant annotations. By focusing exclusively on PPGL and incorporating detailed genotypic and phenotypic annotations, PPGLomics v1.0 enables analyses that would otherwise require substantial bioinformatics expertise. The current version of PPGLomics v1.0 focuses on transcriptomics data from two well-characterized cohorts. Future directions will expand the platform to incorporate additional multi-omics data, including proteomics, metabolomics, immunomics, and single-cell RNA sequencing data as these become available. By integrating transcriptomic data with disease-specific molecular, clinical and other annotations, PPGLomics v1.0 provides a structured foundation for advanced downstream analyses. PPGLomics v1.0 is freely available at https://alkaissilab.shinyapps.io/PPGLomics/.

## Data availability

The PPGLomics Shiny application and associated code are publicly available at https://github.com/HussamAlkaissi/PPGLomics.

## Notes

**Funding** This research was in part supported by the Intramural Research Program of the National Institute of Diabetes and Digestive and Kidney Diseases (NIDDK), National Institute of Dental and Craniofacial Research (NIDCR) and Eunice Kennedy Shriver National Institute of Child Health and Human Development (NICHD) within the National Institutes of Health (NIH). The contributions of the NIH author(s) are considered Works of the United States Government. The findings and conclusions presented in this paper are those of the author(s) and do not necessarily reflect the views of the NIH or the U.S. Department of Health and Human Services.

https://alkaissilab.shinyapps.io/PPGLomics/

### Competing Interest Statement

The authors have declared no competing interest.

## References

1. Lander ES, Linton LM, Birren B, Nusbaum C, Zody MC, Baldwin J, et al. Initial sequencing and analysis of the human genome. Nature. 2001;409(6822):860–921.

2. Berger B, Yu YW. Navigating bottlenecks and trade-offs in genomic data analysis. Nat Rev Genet. 2023;24(4):235–50.

3. Wojcik MH, Larkin K, Cipicchio M, Doupnik A, Zhao C, Cech C, et al. Toward Same-Day Genome Sequencing in the Critical Care Setting. N Engl J Med. 2025;393(20):2063–5.

4. Cancer Genome Atlas Research N, Weinstein JN, Collisson EA, Mills GB, Shaw KR, Ozenberger BA, et al. The Cancer Genome Atlas Pan-Cancer analysis project. Nat Genet. 2013;45(10):1113–20.

5. Cerami E, Gao J, Dogrusoz U, Gross BE, Sumer SO, Aksoy BA, et al. The cBio cancer genomics portal: an open platform for exploring multidimensional cancer genomics data. Cancer Discov. 2012;2(5):401–4.

6. Tang Z, Kang B, Li C, Chen T, Zhang Z. GEPIA2: an enhanced web server for large-scale expression profiling and interactive analysis. Nucleic Acids Res. 2019;47(W1):W556-W60.

7. Chandrashekar DS, Bashel B, Balasubramanya SAH, Creighton CJ, Ponce-Rodriguez I, Chakravarthi B, Varambally S. UALCAN: A Portal for Facilitating Tumor Subgroup Gene Expression and Survival Analyses. Neoplasia. 2017;19(8):649–58.

8. Tang G, Cho M, Wang X. OncoDB: an interactive online database for analysis of gene expression and viral infection in cancer. Nucleic Acids Res. 2022;50(D1):D1334-D9.

9. Goldman MJ, Craft B, Hastie M, Repecka K, McDade F, Kamath A, et al. Visualizing and interpreting cancer genomics data via the Xena platform. Nat Biotechnol. 2020;38(6):675–8.

10. Huang J, Mao L, Lei Q, Guo AY. Bioinformatics tools and resources for cancer and application. Chin Med J (Engl). 2024;137(17):2052–64.

11. Richter S, Qiu B, Ghering M, Kunath C, Constantinescu G, Luths C, et al. Head/neck paragangliomas: focus on tumor location, mutational status and plasma methoxytyramine. Endocr Relat Cancer. 2022;29(4):213–24.

12. Hamidi O, Young WF, Jr., Iniguez-Ariza NM, Kittah NE, Gruber L, Bancos C, et al. Malignant Pheochromocytoma and Paraganglioma: 272 Patients Over 55 Years. J Clin Endocrinol Metab. 2017;102(9):3296–305.

13. Alkaissi H, Taieb D, Lin FI, Del Rivero J, Wang K, Clifton-Bligh R, Pacak K. Approach to the Patient with Metastatic Pheochromocytoma and Paraganglioma. J Clin Endocrinol Metab. 2025.

14. Mete O, Asa SL, Gill AJ, Kimura N, de Krijger RR, Tischler A. Overview of the 2022 WHO Classification of Paragangliomas and Pheochromocytomas. Endocr Pathol. 2022;33(1):90–114.

15. Pacak K. New Biology of Pheochromocytoma and Paraganglioma. Endocr Pract. 2022;28(12):1253–69.

16. Fishbein L, Leshchiner I, Walter V, Danilova L, Robertson AG, Johnson AR, et al. Comprehensive Molecular Characterization of Pheochromocytoma and Paraganglioma. Cancer Cell. 2017;31(2):181–93.

17. Dahia PL. Pheochromocytoma and paraganglioma pathogenesis: learning from genetic heterogeneity. Nat Rev Cancer. 2014;14(2):108–19.

18. Cascon A, Calsina B, Monteagudo M, Mellid S, Diaz-Talavera A, Curras-Freixes M, Robledo M. Genetic bases of pheochromocytoma and paraganglioma. J Mol Endocrinol. 2023;70(3).

19. Flynn A, Pattison AD, Balachander S, Boehm E, Bowen B, Dwight T, et al. Multi-omic analysis of SDHB-deficient pheochromocytomas and paragangliomas identifies metastasis and treatment-related molecular profiles. Nat Commun. 2025;16(1):2632.

20. Flynn A, Pattison AD, Balachander S, Boehm E, Bowen B, Dwight T, et al. Multi-omic analysis of SDHB-deficient pheochromocytomas and paragangliomas identifies metastasis and treatment-related molecular profiles. Res Sq. 2024.

21. Backman S, Botling J, Nord H, Ghosal S, Stalberg P, Juhlin CC, et al. The evolutionary history of metastatic pancreatic neuroendocrine tumours reveals a therapy driven route to high-grade transformation. J Pathol. 2024;264(4):357–70.

22. Serafim RB, Cardoso C, Arfelli VC, Valente V, Archangelo LF. PIMREG expression level predicts glioblastoma patient survival and affects temozolomide resistance and DNA damage response. Biochim Biophys Acta Mol Basis Dis. 2022;1868(6):166382.

23. Jiang L, Ren L, Zhang X, Chen H, Chen X, Lin C, et al. Overexpression of PIMREG promotes breast cancer aggressiveness via constitutive activation of NF-kappaB signaling. EBioMedicine. 2019;43:188–200.

24. Bolanos-Garcia VM, Blundell TL. BUB1 and BUBR1: multifaceted kinases of the cell cycle. Trends Biochem Sci. 2011;36(3):141–50.

25. Malumbres M. Physiological relevance of cell cycle kinases. Physiol Rev. 2011;91(3):973–1007.

26. Ciciro Y, Ragusa D, Sala A. Expression of the checkpoint kinase BUB1 is a predictor of response to cancer therapies. Sci Rep. 2024;14(1):4461.

27. Zheng D, Li J, Yan H, Zhang G, Li W, Chu E, Wei N. Emerging roles of Aurora-A kinase in cancer therapy resistance. Acta Pharm Sin B. 2023;13(7):2826–43.

